# Inducible expression of immediate early genes is regulated through dynamic chromatin association by NF45/*ILF2* and NF90/*ILF3*

**DOI:** 10.1101/261305

**Authors:** Ting-Hsuan Wu, Lingfang Shi, Anson W. Lowe, Mark R. Nicolls, Peter N. Kao

## Abstract

Immediate early gene (IEG) transcription is rapidly activated by diverse stimuli without requiring new protein synthesis. This transcriptional regulation is assumed to involve constitutively expressed nuclear factors that are targets of signaling cascades initiated at the cell membrane. NF45 and its heterodimeric partner NF90 are chromatin-interacting proteins that are constitutively expressed and localized predominantly in the nucleus. Previously, NF90 chromatin immunoprecipitation followed by deep sequencing (ChIP-seq) in K562 erythroleukemia cells revealed its enriched association with chromatin at active promoters and strong enhancers. NF90 specifically occupied the promoters of IEGs. Here, ChIP in serum-starved HEK293 cells demonstrated that NF45 and NF90 pre-exist and specifically occupy the promoters of IEG transcription factors *EGR1*, *FOS* and *JUN*. Cellular stimulation with phorbol myristyl acetate increased NF90 occupancy, while decreasing NF45 occupancy at promoters of *EGR1*, *FOS* and *JUN*. In HEK293 cells stably transfected with doxycycline-inducible shRNA vectors targeting NF90 or NF45, doxycycline-mediated knockdown of NF90 or NF45 attenuated the inducible expression of *EGR1*, *FOS*, and *JUN* at the levels of mRNA and protein. NF90 and NF45 operate as constitutively-expressed transcriptional regulators of IEGs. Dynamic chromatin association of NF45 and NF90 at IEG promoters are observed upon stimulation, and NF45 and NF90 contribute to inducible expression of IEGs. NF45 and NF90 operate as chromatin regulators of the immediate early response.

## INTRODUCTION

The rapid cellular response that occurs upon recognition of biological or environmental signals is crucial for adaptation and survival of the organism (1-3). The subset of genes that are rapidly expressed upon induction are termed immediate early genes (IEGs) (4). Inducible expression of IEGs in response to diverse regulatory signals underlies acute inflammation (5-7), neuronal activity (8), cell proliferation, and differentiation (1,9,10). Aberrant expression of IEGs is involved in malignant cellular transformation (11) and is a feature of diverse cancers (12,13).

Upon stimulation, initial expression of IEGs occurs on the timescale of minutes to hours (4,14). The protein products of these IEGs critically include transcription factors such as *EGR1*, *FOS*, and *JUN*, which then activate and induce expression of secondary response genes (14). Regulation of this hierarchical program upon cellular stimulation does not require *de novo* protein synthesis. Transcriptional regulation of IEGs is thus assumed to involve pre-existing nuclear factors that are constitutively expressed, which are targets of signaling cascades initiated at the cell membrane.

Features of IEG promoters include over-representation of transcription factor binding sites and high affinity TATA boxes (4). Chromatin structure of IEGs shows enrichment of active chromatin marks and “poised” accumulation of RNA polymerase II (14). Stimulation-induced chromatin remodeling at promoters of IEGs exposes specific DNA binding sequences for transcription factors such as serum-response factor (SRF), nuclear factor kappa B (NF-κB), and cyclic AMP response element-binding protein (CREB) (4). Transcription of DNA by RNA Polymerase II complex into RNA is followed by post-transcriptional regulation at the levels of RNA splicing, nuclear export, stabilization, and translational regulation of the nascent transcripts (4).

Nuclear Factor 90 (NF90, encoded by the *ILF3* gene) and Nuclear Factor 45 (NF45, encoded by the *ILF2* gene) are multifunctional DNA- and RNA-binding proteins originally purified and cloned based on their inducible and specific DNA-binding to the NF-AT/ ARRE-2 sequence in the *IL2* promoter from activated Jurkat T-cells (15,16). NF90 and NF45 interact tightly as a heterodimer through their shared dimerization zinc-finger (DZF) domains (17). NF90 has two dsRNA binding domains, and both NF90 and NF45 contain an arginine/glycine/glycine (RGG) domain that is capable of binding to nucleic acids (18). The interactions of NF90 and NF45 with chromatin have been demonstrated at several regulatory regions in addition to *IL2* (19-21), including promoters of *FOS* (22), *PLAU* (23)and enhancer of HLA-DR alpha (24) and *IL13* (25).

NF90 and NF45 have been shown to regulate embryonic pluripotency, and development. NF90 is required for normal development. Mice with targeted disruption of NF90 were born small and weak and succumbed to perinatal death from neuromuscular respiratory failure (19). NF45 knockout in mice resulted in early embryonic lethality^1^. NF45 physically interacts with Oct4 and Nanog in embryonic stem cells (ESC) to promote pluripotency (26). Targeted disruption of NF90 and NF45 impaired ESC proliferation and promoted differentiation to an epiblast-like state (27). NF90 and NF45 regulate cell cycle progression (18,19,28), cell growth and proliferation (28-33), and are amplified, overexpressed and mutated in diverse cancers (34,35).

We recently characterized NF90 as a transcription factor involved in promoting proliferation and renewal over differentiation in K562 erythroleukemia cells using chromatin immunoprecipitation followed by deep sequencing (ChIP-seq) (36). Rigorous statistical testing between biological replicates with Irreproducible Discovery Rate (IDR) analysis revealed chromatin occupancy of NF90 at 9,081 specific genomic sites, with over a third occurring at promoters of protein-coding genes.

Further analysis of NF90 chromatin occupancy in a context of histone modifications revealed enrichment of NF90 occupancy frequencies at active promoters and strong enhancers. To investigate the functional role of NF90 in transcriptional regulation in K562 cells, we compared the 2,927 genes with NF90 chromatin occupancy in its proximal promoter to a dataset of 446 genes that were differentially expressed upon NF90 knockdown by shRNA. In K562 cells grown in 10% serum at basal growth conditions, this integrated analysis of genes under transcription regulation by NF90 revealed an overrepresentation of IEGs.

Tullai *et al*. previously demonstrated that the cellular response to growth factor stimulation involves initial induction of IEGs, followed by delayed expression of primary response genes that also do not require initial protein synthesis, and finally, secondary response genes (14). Here, we examined NF90 ChIP-seq data in K562 cells at normal growth conditions. Compared to delayed primary response genes or secondary response genes, we found enriched NF90 occupancy at promoters of IEGs, including transcription factors *EGR1*, *FOS*, and *JUN*.

The chromatin occupancy of NF90 at promoters of IEGs in normally growing K562 cells suggested to us that NF90, together with its heterodimeric partner NF45, might contribute to regulation of IEG expression. In this study, we test the hypothesis that NF90 and NF45 contribute to transcriptional regulation of IEG transcription factors *EGR1*, *FOS*, and *JUN*. In HEK293 cells that were serum starved, we stimulated with phorbol 12-myristate 13-acetate (PMA) to induce expression of IEGs and examine the early recruitment of NF90 and NF45 to promoters of immediate early transcription factors using ChIP.

In HEK293 cells, serum starvation followed by stimulation with phorbol 12-myristate 13-acetate (PMA) induces dynamic association of NF90 and NF45 at promoters of immediate early transcription factors *EGR1*, *FOS*, and *JUN*. In HEK293 cells stably transfected with doxycycline-inducible shRNA vectors targeting NF90 or NF45, we demonstrate that doxycycline-mediated knockdown of NF90 or NF45 attenuated the PMA-inducible expression of *EGR1*, *FOS*, and *JUN* at levels of RNA and protein.

Taken together, we present evidence that NF90 and NF45 are constitutively expressed chromatin-interacting proteins pre-existing at the promoters of IEGs. Dynamic association of NF90 and NF45 at proximal promoters is observed upon cell stimulation, and NF90 and NF45 proteins each contribute positively to inducible expression of IEGs. NF90 and NF45 operate as chromatin regulators that activate the immediate early response.

## EXPERIMENTAL PROCEDURES

### Cell culture and stimulation

Human embryonic kidney (HEK) 293 cells (ATCC, Manassas, VA) were maintained in Dulbecco′s modified Eagle′s medium supplemented with 10% fetal calf serum with Penicillin and Streptomycin. Cells were serum starved for 12 h before stimulation with 20 ng/ml phorbol 12-myristate 13-acetate (PMA) for indicated durations, harvested by Trypsin-EDTA, centrifugation, and frozen at - 80°C.

### Reagents

The following mouse monoclonal antibodies were used for chromatin immunoprecipitation and Western immunoblotting: anti-NF90 (mouse mAb DRBP76; BD 612154), anti-NF45 (mouse mAb NF45 H-4; Santa Cruz 365283). Rabbit monoclonal antibodies used for Western immunoblotting include: anti-EGR1 (rabbit mAb EGR1 15F7; Cell signaling 4153), anti-FOS (rabbit mAb FOS 9F6; Cell signaling 2250), anti-JUN (rabbit mAb JUN 60A8; Cell signaling 9165).

### Chromatin immunoprecipitation

Cells were harvested and crosslinked in 1% formaldehyde for 10 minutes at room temperature. Pellets were re-suspended in hypotonic buffer (20 mM HEPES, 10 mM KCl, 1mM EDTA, 10% glycerol) supplemented with Pierce Protease Inhibitor Tablets (Thermo Fisher), placed on ice for 10 minutes and centrifuged to remove cytoplasmic supernatant. Chromatin was released from nuclear pellet with mild chemical lysis with Radio-immunoprecipitation assay buffer (RIPA) containing 0.1% SDS and 1% Triton-X-100. Protein-bound chromatin was sheared with mechanical sonication (Branson 250 Sonifier) at 35% output with 20 second pulses for 30 cycles to obtain chromatin fragments of resolution at 200 – 500 bps. Successful chromatin fragmentation was assayed by reverse cross-linking and electrophoresis through 0.8% agarose gel to confirm enrichment of small chromatin fragments at the target range. Chromatin fragments were pre-cleared with untreated protein A/G agarose beads (Santa Cruz), then incubated in protein A/G agarose beads that were pre-conjugated with monoclonal antibodies to NF90 (BD mAB DRBP76) or NF45 (Santa Cruz sc-365283) overnight at 4°C. Chromatin fragments enriched for NF90 or NF45-binding were eluted in 1% SDS at 65°C for 15 minutes. Chromatin was reverse cross-linked from protein at 65°C overnight. RNase A (100 ug, Sigma Aldrich) was added to sample to incubate at 37°C for 30 minutes, and Proteinase K (100 ug, Invitrogen) was then added to sample to incubate at 45°C for 30 minutes. ChIP DNA was purified using PCR purification column (Qiagen) per manufacturer protocol and eluted in 35 ul EB buffer (10 mM Tris-Cl, pH 8.5).

ChIP DNA was used as template for polymerase chain reaction using primers amplifying specific genomic regions to interrogate NF90 or NF45 binding. PCR primers were synthesized by the Protein and Nucleic Acid Facility at Stanford University, and designed to amplify the *EGR1* proximal promoter: F 5’ – TTCCCCAGCCTAGTTCACGCCTAGGAGCC – 3’, R 5’ – ATATGGCATTTCCGGGTCGCAGCTGG – 3’; *FOS* proximal promoter: F 5’ – CCGCGAGCAGTTCCCGTCAATCCCTC – 3’, R 5’ – GCAGTTCCTGTCTCAGAGGTCTCGTGGGC – 3’; *JUN* proximal promoter: F 5’ –CCTCCCGGGTCCCTGCATCCCC – 3’, R 5’ – ACGCCTCTCGGCCCTCTCTTCCC – 3’; *HBB* locus: F 5’ – CCTCGGCCTCCCAAAGTGCCAGGATTACAG – 3’, R 5’ – ACAAGCATGCGTCACCATGCCTGGC – 3’. PCR amplifications were performed using a three-step protocol with 59°C annealing temperature and 35 cycles.

### Molecular cloning and transfection

The pINDUCER lentiviral toolkit for inducible RNA interference (Meerbrey *et al*., 2011) was utilized for inducible shRNA knockdown of NF90 or NF45. pInducer10-mir-RUP-PheS was a gift from Stephen Elledge (Addgene plasmid # 44011), containing a constitutively expressed transcript encoding the puromycin resistance gene, and a doxycycline-inducible cassette that expresses target shRNA sequence as well as turboRFP fluorescent protein. Validated shRNA sequences D2 directed against NF90 (5’ – GUGCUGGUUCCAACAAAA – 3’) and D5 directed against NF45 (5’ – AGUCGUGGAAAGCCUAAGA – 3’) were designed as described by Guan *et al*. (19) and sub-cloned into the doxycycline-inducible cassette driven by the tetracycline-responsive TRE2 promoter in pINDUCER10 to create plasmids pINDUCER10-shNF90 (D2) and pINDUCER10-shNF45 (D5).

HEK293 cells at 70% confluency were transfected with either pINDUCER10-shNF90 (D2) or pINDUCER10-shNF45 (D5) using JetPrime transfection reagent (Polyplus). Beginning at 48 h after transfection, drug selection for stable transfection was imposed using puromycin (1 μg/ml).

HEK293 cells stably expressing D2 or D5 were treated without or with doxycycline (1 ug/ul) in DMEM medium supplemented with 10% FBS for 96 h to attenuate NF90 or NF45 expression. Cells were then serum-starved for 12 h with continuing presence of doxycycline, before stimulation with PMA for the indicated durations.

### Reverse transcription polymerase chain reaction

RNA levels were assayed using reverse transcription PCR. Total RNA was isolated from cells using the RNeasy mini kit (Qiagen). First-strand cDNA was synthesized from 2 ug of total RNA using Superscript IV VILO reverse transcriptase (Invitrogen) with a mix of oligo(dT) and random hexamer primers per manufacturer protocol, which was used as template for polymerase chain reactions. Primers were designed to amplify: *EGR1*: F 5’ – GCACCTGACCGCAGAGTCTTTTCCT – 3’, R 5’ – GGTGTTGCCACTGTTGGGTGCAG –3’; *FOS*: F 5’ – CTGTCAACGCGCAGGACTTCTGC – 3’, R 5’ – GCTCGGCCTCCTGTCATGGTCT – 3’; *JUN*: F 5’ – CGATGCCCTCAACGCCTCGTTC – 3’, R 5’ – GTGATGTGCCCGTTGCTGGACTG– 3’; *ACTB*: F 5’ – CCAATCAGCGTGCGCCGTTCC– 3’, R 5’ – ATCATCCATGGTGAGCTGGCGG– 3’. PCR amplifications were performed using a three-step protocol with 55°C annealing temperature and 30 cycles.

### Western immunoblotting

Cells were harvested and whole cell extracts were prepared by incubation on ice for 30 minutes in 8M urea lysis buffer (8M urea, 300 mM NaCl, 0.5% NP 40, 50 mM Na_2_HPO_4_, 50 mM Tris-HCl, 1 mM PMSF) supplemented with Pierce Protease Inhibitor Tablets (Thermo Fisher). Total protein of 20 ug was separated on 7.5% or 10% SDS-PAGE using Mini-Protean II apparatus (Bio-Rad) and transferred to polyvinylidene difluoride (PVDF) membranes. Membranes were blocked in 5% nonfat dry milk in TBST, probed with indicated antibodies at 1:1000 dilution in 5% milk overnight at 4°C, washed three times in TBST, incubated with anti-mouse or anti-rabbit horseradish peroxidase-coupled secondary antibody (Santa Cruz) at 1:10,000 dilution in 5% milk for 1 h at room temperature, and visualized with enhanced chemiluminescence detection (Amersham).

### Immunofluorescence staining

Cells were passaged and seeded on chamber slides. Upon serum starvation and stimulation, cells were fixed and permeabilized in 100% methanol at room temperature for 15 minutes, and bleaching of native fluorescence was confirmed with microscopy. After rinsing three times with PBS, cells were blocked in 5% FBS/ 0.3% Triton X-100 in PBS for 1 h at room temperature. The chamber slides were then incubated with anti-NF90 (mouse mAb BD, 1:200) or anti-NF45 (mouse mAb Santa Cruz, 1:200); and anti-EGR1 (rabbit mAb Cell Signaling, 1:100), anti-FOS (rabbit mAb Cell Signaling, 1:100), or anti-JUN (rabbit mAb Cell Signaling, 1:100) diluted in 0.5% FBS/ 0.03% Triton X-100 in PBS overnight at 4°C. After overnight incubation, unbound primary antibody was removed by washing the slides three times in PBS. Fluorescent conjugated secondary antibodies anti-mouse (Goat anti-Mouse IgG Alexa Fluor 488 Invitrogen, 1:500) and anti-rabbit (Goat anti-Rabbit IgG Alexa Fluor 594 Invitrogen, 1:500) were applied to slides for 1 h at room temperature protected from light. Slides were washed three times with PBS, counterstained with DAPI nuclear stain (Pierce, 1 mg/ml), and mounted using VECTASHIELD antifade solution (Vector Laboratories). Imaging was performed on a Zeiss confocal laser scanning microscope (LSM 880, Zeiss) with a 20X objective lens.

## RESULTS

### Enriched NF90 chromatin occupancy at promoters of immediate early genes

We recently used ChIP-seq to demonstrate the wide chromatin occupancy of NF90 {Wu, 2017 #1840}. Here, we further characterized NF90 chromatin occupancy in the growing K562 erythroleukemia cell line to reveal selective enrichment at promoters of IEGs, including the transcription factors *EGR1*, *FOS*, and *JUN*, known regulators of inflammation, proliferation, and neuronal activity.

The cellular response to growth stimulation involves a sequential gene activation program initiated by the immediate early induction of transcription factors, followed by expression of delayed primary response genes, followed by protein synthesis-dependent expression of secondary response genes (14). We discovered enriched NF90 chromatin occupancy in the promoters of IEGs. The average chromatin occupancy profile of NF90 is presented for the proximal promoters of IEGs, delayed primary response genes (D-PRG), and secondary response genes (SRG) (**Figure 1a**). The specific genes in each group and the genomic coordinates of the proximal promoters are tabulated (**Supplementary Table**). NF90 chromatin occupancy frequency was highest at IEGs, followed by D-PRGs, then SRGs. There was approximately a two-fold enrichment in NF90 occupancy frequency at IEGs compared to SRGs.

**Figure 1.**
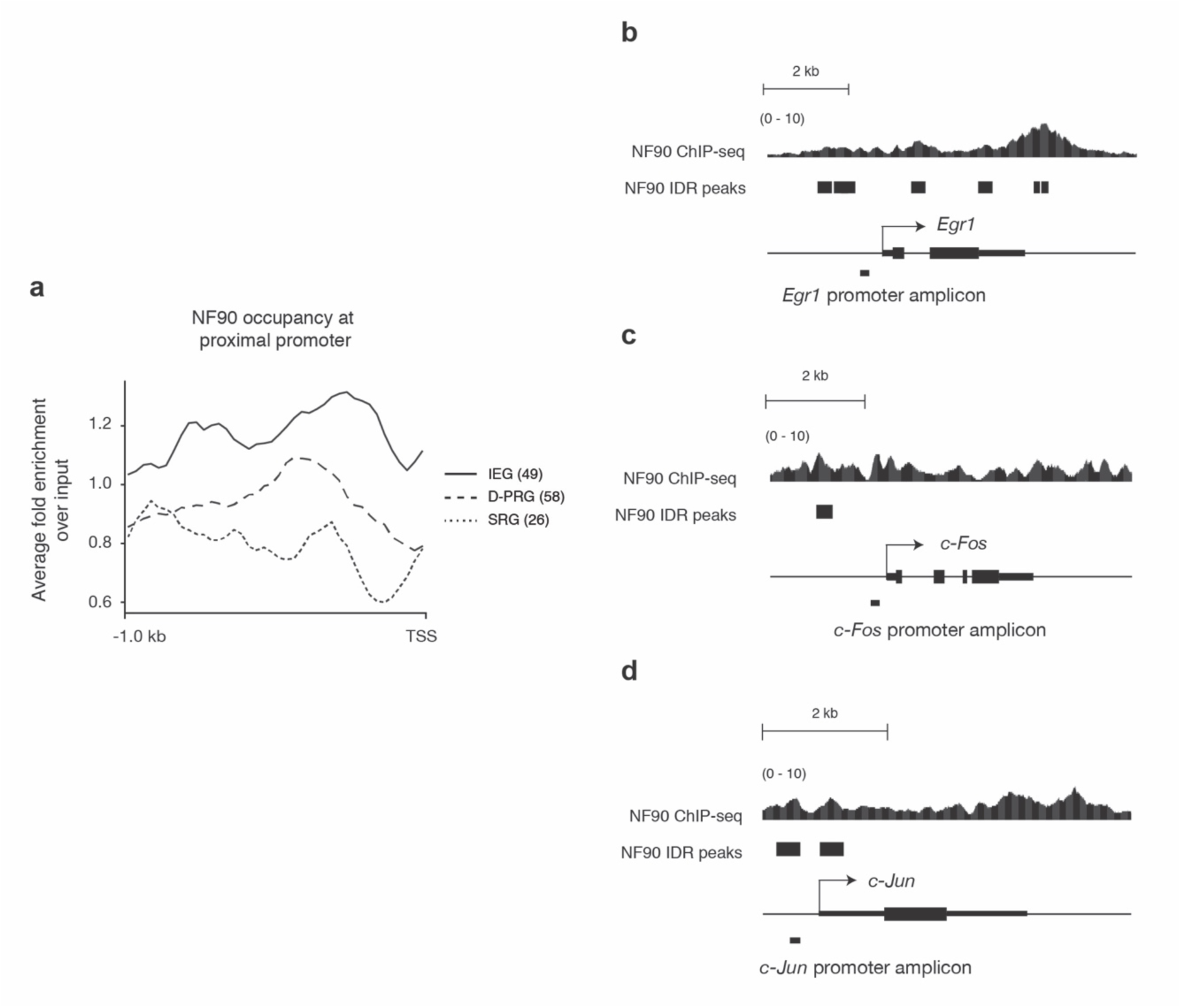
NF90 occupancy at promoters of immediate early genes in K562 cells. **a.** Average occupancy profile of NF90 at proximal promoters within 1000 bp upstream of transcription start site (TSS) of 49 immediate early genes (IEG), 58 delayed primary response genes (D-PRG), and 26 secondary response gene (SRG) defined by Tullai et al. x-axis: relative position near TSS. y-axis: fold-change over input of NF90 ChIP-Seq signal. **b-d.** Signal tracks of NF90 chromatin immunoprecipitation deep sequencing (ChIP-seq) occupancy at promoters of IEGs *EGR1*, *FOS*, and *JUN*; retrieved from UCSC genome browser. K562 cells cultured under basal growth conditions (10% serum) were subject to NF90 ChIP-seq according to standard ENCODE protocols and uniform processing pipelines using antibodies against NF90 (mAb DRBP76; BD). NF90 ChIP-seq signal (fold change over input) is presented, aligned to peaks called by Irreproducible Discovery Rate (IDR) analysis that measures consistency between replicates. Relative position of the amplicon in the proximal promoter of each gene that are used for subsequent validation ChIP experiments are indicated.

We previously proposed that NF90 hierarchically regulates transcription factors to promote cell growth and proliferation over differentiation. Here, we focused on NF90 chromatin occupancy at the promoters of IEGs that are transcription factors. We sought to determine whether NF90, and its heterodimeric partner NF45, might hierarchically regulate transcription factors that control the immediate early responses of cells to diverse stimuli. NF90 chromatin occupancy frequency (fold-change over input) is graphed as a continuous variable, and Irreproducible Discovery Rate (IDR) analysis marks the discrete genomic intervals where rigorous statistical testing between biological replicates indicates strong confidence in NF90 chromatin occupancy (**Figure 1b-d**). At the loci for *EGR1*, *FOS*, and *JUN,* continuous NF90 chromatin occupancy is found at the transcription start site (TSS), throughout the gene body, and at the transcription end site (TES), consistent with previous findings. Most significantly, we demonstrate NF90 chromatin occupancy at proximal promoters of immediate early transcription factors *EGR1*, *FOS*, and *JUN* (**Figure 1b-d**).

### Dynamic chromatin association of NF90 and NF45 at promoters of immediate early genes during cell stimulation

To confirm and generalize our finding that NF90 occupied the promoters of immediate early transcription factors in normal growing K562 cells, we extended our study of IEG regulation to the human embryonic kidney (HEK) 293 cell line, which demonstrates no tissue-specific gene expression signature and a transformed immortal phenotype (37).

Based on the bioinformatics results that demonstrated enrichment of NF90 occupancy at promoters at immediate early transcription factors, and the frequent dimerization interaction between NF90 and NF45 mediated by their shared domain associated with zinc fingers (DZF), we hypothesized that, upon cellular stimulation, NF90 and NF45 may coordinately regulate expression of immediate early transcription factors *EGR1*, *FOS*, and *JUN*. The stimulation of choice for these experiments was phorbol 12-myristate 13-acetate (PMA), an activator of protein kinase C (PKC), previously demonstrated to stimulate inducible binding of NF90 to the *IL2* promoter (20, 21), and known to rapidly induce expression of IEGs, including *EGR1*, *FOS*, and *JUN* (40). For maximal contrast of IEG induction, HEK293 cells were serum starved overnight. Quiescent cells were then stimulated with 20 ng/ml PMA to achieve rapid induction of IEGs.

To test the hypothesis that NF90 and NF45 may exhibit dynamic association to the promoters of immediate early transcription factors, we performed ChIP using monoclonal antibodies to NF90 or NF45, and designed PCR primers to interrogate the proximal promoter regions within 500 bp upstream of the transcription start sites of *EGR1*, *FOS*, and *JUN* that demonstrated high NF90 chromatin occupancy frequency in K562 cells (**Figure 1b-d**).

We characterized chromatin occupancy of NF90 and NF45 to the *EGR1*, *FOS*, and *JUN* promoters in quiescent and stimulated HEK293 cells *in vivo*, using formaldehyde crosslinking and chromatin immunoprecipitation with antibodies against NF90 or NF45 (**Figure 2**). We examined the transcription factor binding tracks deposited on ENCODE and used a region in the beta globin (*HBB*) locus as a negative control for its lack of transcription factor binding and low transcriptional activity in HEK293 cells. PCR using input chromatin prepared from quiescent and stimulated HEK293 cells as template resulted in amplification products by primers to the *EGR1*, *FOS*, and *JUN* promoters, and the *HBB* locus, and these are constant across 0, 30, and 60 minutes of PMA stimulation (**Figure 2a,c,** lanes 1-3).

**Figure 2.**
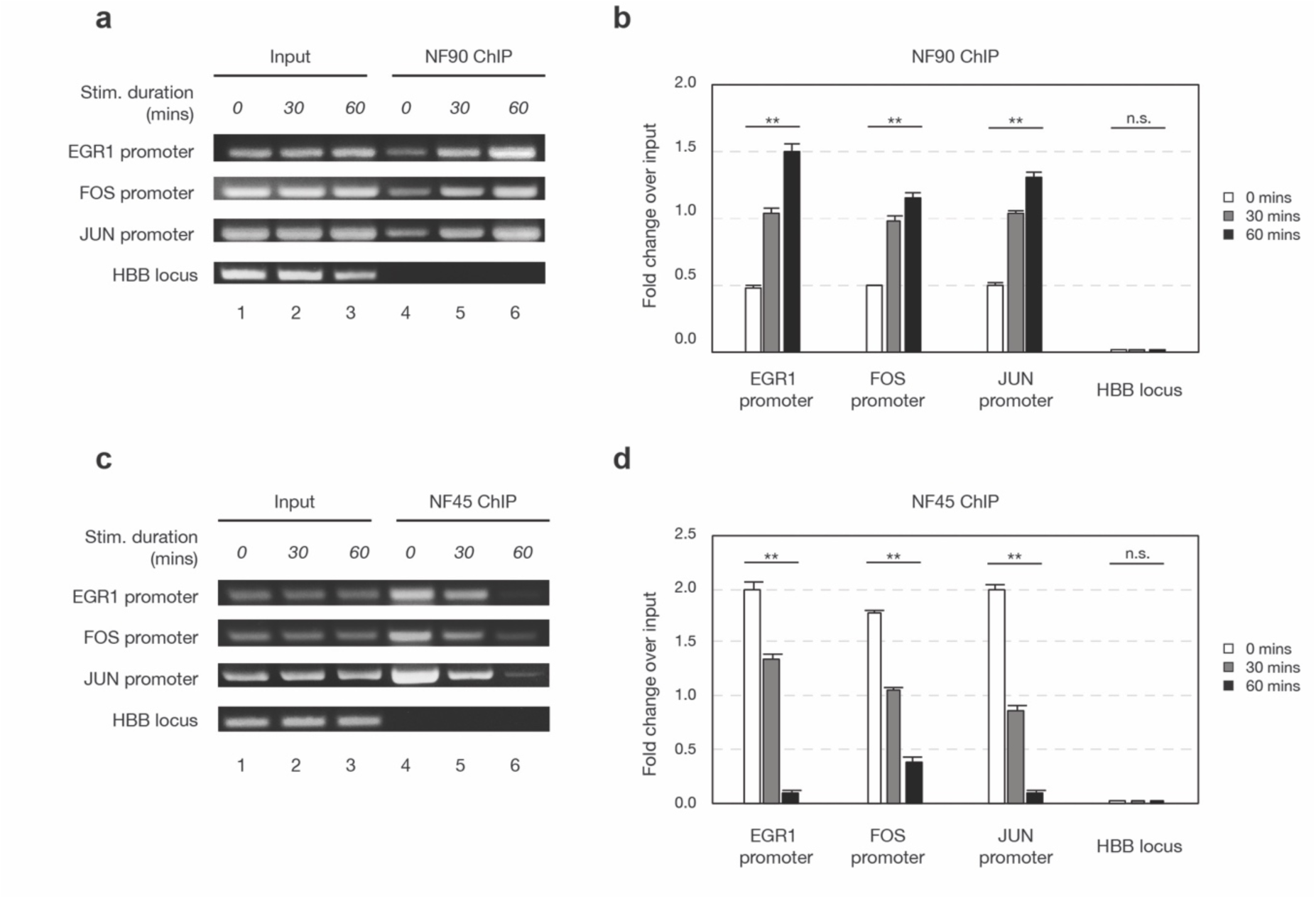
Dynamic and reciprocal association of NF90 and NF45 to promoters of IEGs upon stimulation. HEK293 cells were serum starved for 24 h, then stimulated by PMA (20 ng/ml) for indicated durations. Chromatin immunoprecipitation was performed with antibodies against NF90 (**a-b**) or NF45 (**c-d**). Presence and abundance of IEG promoter fragments were assessed with polymerase chain reaction subject to 1.5% agarose electrophoresis. **a**, **c**, Representative images of ChIP-PCR experiments. **b**, **d**, ImageJ semi-quantification of **a** and **c** (mean ± SEM, *n* = 3). One-way analysis of variance (ANOVA) test, ** p < 0.01.

Immunoprecipitation from input chromatin using antibody against NF90 showed no precipitation of the *HBB* locus across time points (**Figure 2a**, lanes 4-6). NF90 occupancy at proximal promoters of *EGR1*, *FOS*, and *JUN* in nonstimulated HEK293 cells was present but modest (**Figure 2a, lane 4**). Upon PMA stimulation, we observed a substantial increase of NF90 chromatin occupancy at the proximal promoters of *EGR1*, *FOS*, and *JUN* at 30 minutes, and a further increase at 60 minutes (**Figure 2a**, lanes 5,6 vs. 4; quantified in **Figure 2b**).

Similarly, immunoprecipitation from input chromatin using antibody against NF45 showed no precipitation of the *HBB* locus across time points (**Figure 2c**, lanes 4-6). In contrast with NF90 ChIP, we observed strong NF45 chromatin occupancy at proximal promoters of *EGR1*, *FOS*, and *JUN* in nonstimulated HEK293 cells (**Figure 2c**, lane 4). Remarkably, upon PMA stimulation, we observed a substantial decrease of NF45 chromatin occupancy at the proximal promoters of *EGR1*, *FOS*, and *JUN* at 30 minutes (**Figure 2c**, lane 5 vs. 4), and further decrease of NF45 chromatin occupancy at 60 minutes (**Figure 2c**, lane 6 vs. 4; quantified in **Figure 2d**).

This dynamic and reciprocal chromatin association by NF90 and NF45 suggested that, in contrast to frequent observations of NF90 and NF45 as a heterodimer, they are each capable of independent interactions at proximal promoters of immediate early transcription factors. These dynamic chromatin interactions of NF90 and NF45 are likely to be triggered by signaling events initiated at the cell membrane.

### Establishment and characterization of stable HEK293 cells with doxycycline-regulated RNAi targeting NF90 and NF45

The dynamic and reciprocal occupancy of NF90 and NF45 at the proximal promoters of immediate early transcription factors suggested that NF90 and NF45 may coordinately regulate gene activation of transcription factors in the immediate early response. To study the functional role of NF90 and NF45 in regulating immediate early transcription factors *EGR1*, *FOS*, and *JUN*, we used RNA interference using validated (19) shRNA sequences against NF90 (D2) or NF45 (D5) to knockdown NF90 or NF45 proteins and examined the consequences upon inducible expression of *EGR1*, *FOS*, and *JUN*.

Because NF90 and NF45 are genes essential for normal mammalian development, as well as for cellular growth and proliferation, we anticipated that constitutive knockdown in HEK293 cells might adversely affect cell viability, as previously reported (18). Therefore, we sought to establish a system for doxycycline-regulated RNAi knockdown of NF90 and NF45. The pINDUCER vectors are multicistronic plasmids in which a strong ubiquitin promoter drives constitutive expression of reverse tetracycline transactivator protein and resistance to puromycin, and a tetracycline-regulated promoter drives expression of turboRed fluorescent protein and shRNA (41). pINDUCER 10 plasmids were created with shRNA sequences against NF90 (D2) or NF45 (D5). We transfected HEK293 cells with either pINDUCER-shNF90 (D2) or pINDUCER-shNF45 (D5), and applied puromycin selection to generate stably-transfected HEK293 cells with doxycycline-regulated shRNAs directed to NF90 (D2) or NF45 (D5).

The efficacy of doxycycline-regulated knockdown of NF90 or NF45 in stably-transfected HEK293 D2 and D5 cells was characterized by Western immunoblotting (**Figure 3a**). HEK293 cells were untreated (Dox-), or treated for 96 h with 1 μg/ml doxycycline (Dox+), and whole cell lysates prepared with urea to achieve full extraction of these proteins from chromatin. Compared to Dox-, Dox+ HEK293 D2 cells (shRNA targeting NF90) demonstrated substantial reduction in NF90 protein expression (**Figure 3a**, NF90: lanes 3, 4 vs. 1, 2). Additionally, compared to Dox-, Dox+ HEK293 D2 cells showed reduction in NF45 protein expression (**Figure 3a**, NF45: lanes 3,4 vs. 1,2). Compared to Dox-, Dox+ treated HEK293 D5 cells (shRNA targeting NF45) demonstrated substantial reduction in NF45 protein expression (**Figure 3a**, NF45: lanes 7,8 vs. 5,6). Additionally, compared to Dox-, Dox+ HEK293 D5 cells showed reduction in NF90 protein expression (**Figure 3a**, NF90: lanes 7,8 vs. 5,6). Our results demonstrating modest attenuation of the levels of NF45 or NF90 when the heterodimeric partner is targeted by RNAi is consistent with the previous suggestion that NF90 and NF45 interact and confer mutual stabilization (19).

**Figure 3.**
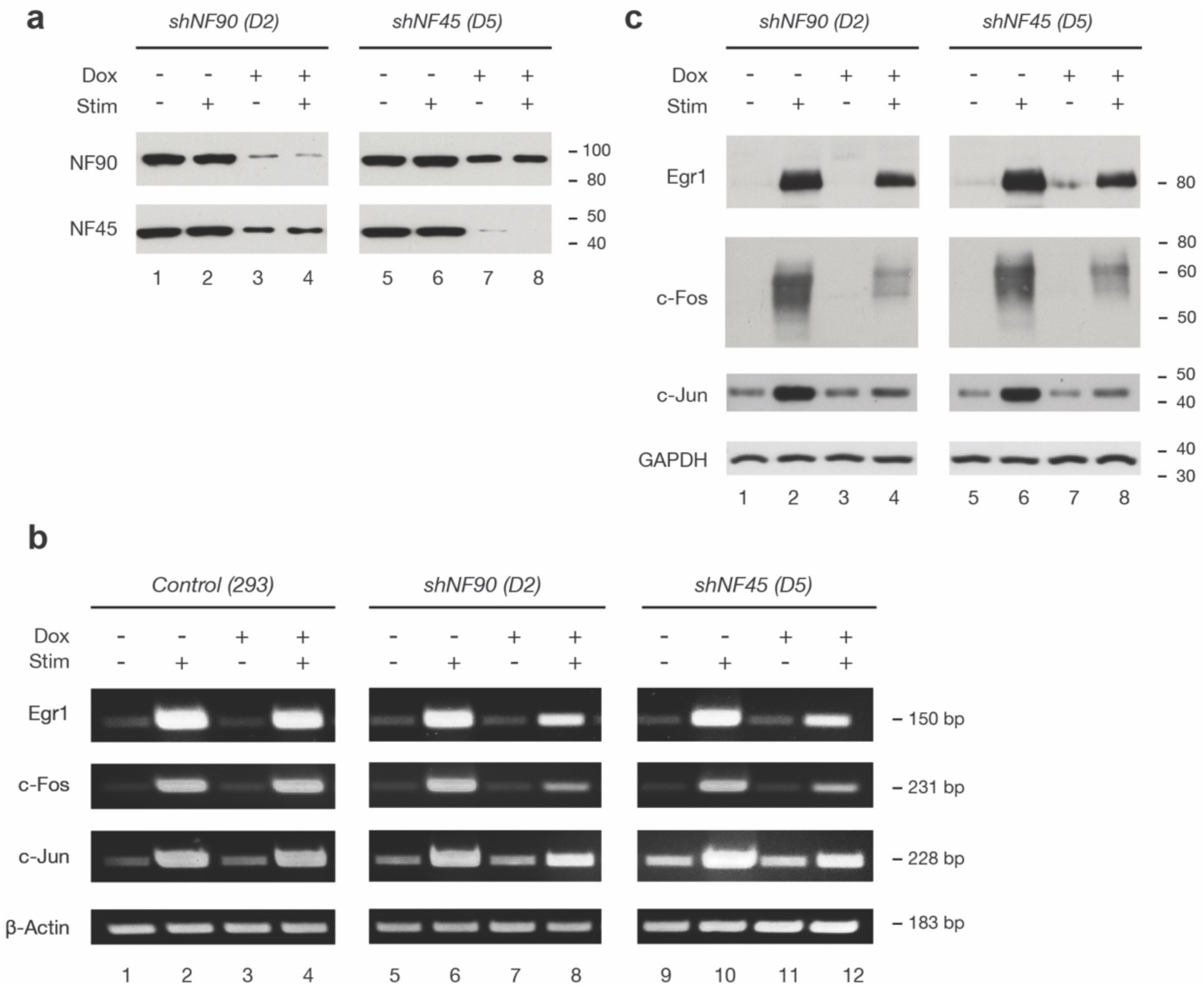
Reduced NF90 or NF45 expression attenuated inducible protein expression of IEGs. HEK293 cells stably transfected with pINDUCER10-shNF90 (D2) or pINDUCER10-shNF45 (D5) were treated without or with doxycycline for 96 h, then serum starved for 12 h and stimulated by PMA (20 ng/ml) for 30 m for strong induction of IEG mRNA, or 2 h for sufficient IEG protein expression. **a**, Doxycycline-mediated knockdown was assessed with western immunoblotting using antibodies against NF90 and NF45. **b**, Reverse transcription polymerase chain reaction was performed on cDNA reverse transcribed from 2 ug total RNA harvested from non-transfected control cells (lanes 1-4), cells stably transfected with shNF90 (D2) (lanes 5-8), or cells stably transfected with shNF45 (D5) (lanes 9-12), using primers specific for transcripts of IEGs *EGR1*, *FOS*, and *JUN*. **c**, Induction of IEGs *EGR1*, *FOS*, and *JUN* were assessed with western immunoblotting.

### NF90 and NF45 contribute to inducible expression of immediate early genes upon cell stimulation

To test the hypothesis that NF90 and NF45 contribute to inducible expression of immediate early transcription factors upon stimulation, we compared the induction of immediate early transcription factors *EGR1*, *FOS*, and *JUN* in HEK293 cells that stably expressed doxycycline-regulated shNF90 (D2) or shNF45 (D5), as well as non-transfected HEK293 cells for control.

In control HEK293 cells that have been serum starved for 12 h to enter quiescence, mRNA expression of *EGR1*, *FOS*, and *JUN* is low to undetectable (**Figure 3b**, lanes 1 and 3). Stimulation of HEK293 cells with PMA (20 ng/ml) for 30 minutes achieved strong induction of mRNA transcripts for immediate early transcription factors *EGR1*, *FOS*, and *JUN* (**Figure 3b**, lane 2 vs 1, lane 4 vs. 3). Beta-actin (*ACTB*) was used as internal control, and *ACTB* mRNA levels were constant across conditions (**Figure 3b**). Compared to Dox-cells, Dox+ HEK293 cells demonstrated no detectable change in the PMA-stimulated induction of *EGR1*, *FOS*, and *JUN* RNAs (**Figure 3b**, lane 4 vs. 2).

We next characterized HEK293 D2 cells stably transfected with doxycycline-regulated shRNA targeting NF90. In Dox-HEK293 D2 cells, the PMA-stimulated RNA levels for *EGR1*, *FOS*, and *JUN* were comparable to those in plain HEK293 cells (**Figure 3b**, lane 6 vs 2). In Dox+ HEK293 D2 cells, the PMA-stimulated RNA levels of *EGR1*, *FOS*, and *JUN* were attenuated compared to Dox-D2 cells (**Figure 3b**, lane 8 vs 6), or Dox+ plain HEK293 cells (**Figure 3b**, lane 8 vs 4).

We next characterized HEK293 D5 cells stably transfected with doxycycline-regulated shRNA targeting NF45. In Dox-HEK293 D5 cells, the PMA-stimulated RNA levels for *EGR1*, *FOS*, and *JUN* were comparable to those in plain HEK293 cells (**Figure 3b**, lane 10 vs 2). In Dox+ HEK293 D5 cells, the PMA-stimulated RNA levels of *EGR1*, *FOS*, and *JUN* were attenuated compared to Dox-D5 cells (**Figure 3b**, lane 12 vs 10), or Dox+ plain HEK293 cells (**Figure 3b**, lane 12 vs 4).

Taken together these results indicate that NF90 and NF45 proteins positively regulate PMA-inducible RNA transcription of immediate early transcription factors *EGR1*, *FOS*, and *JUN*.

To assay the inducible expression of IEGs at the protein level, we stimulated HEK293 cells with PMA for 2 hours and prepared whole cell lysates with urea to quantify by immunoblotting stimulated protein levels of *EGR1*, *FOS*, and *JUN*. We used Glyceraldehyde 3-phosphate dehydrogenase (GAPDH) as an internal control (**Figure 3c**). HEK293 cells stably expressing shNF90 (D2) or shNF45 (D5) were cultured without or with doxycycline for 96 h before serum starvation overnight followed by stimulation with PMA for 2 h. We determined that doxycycline-mediated knockdown of NF90 or NF45 proteins each attenuated the PMA-inducible protein expression of immediate early transcription factor proteins *EGR1*, *FOS*, and *JUN* (**Figure 3c**, lane 4 vs. 2 and lane 8 vs. 6).

We performed immunofluorescence microscopy in HEK293 cells stably expressing doxycycline-regulated shRNA targeting NF90 (D2) or NF45 (D5) to achieve single cell characterization of the effects of NF90 and NF45 protein levels on PMA-inducible expression of immediate early transcription factors, *EGR1*, *FOS*, and *JUN* (**Figure 4,5**). Multiplexed immunofluorescence was achieved using mouse monoclonal antibodies to detect NF90 or NF45 proteins (detected with anti-mouse Alexa Fluor 488 nm secondary antibody), and rabbit monoclonal antibodies to detect *EGR1*, *FOS*, and *JUN* proteins (detected with anti-rabbit Alexa Fluor 594 secondary antibody).

**Figure 4.**
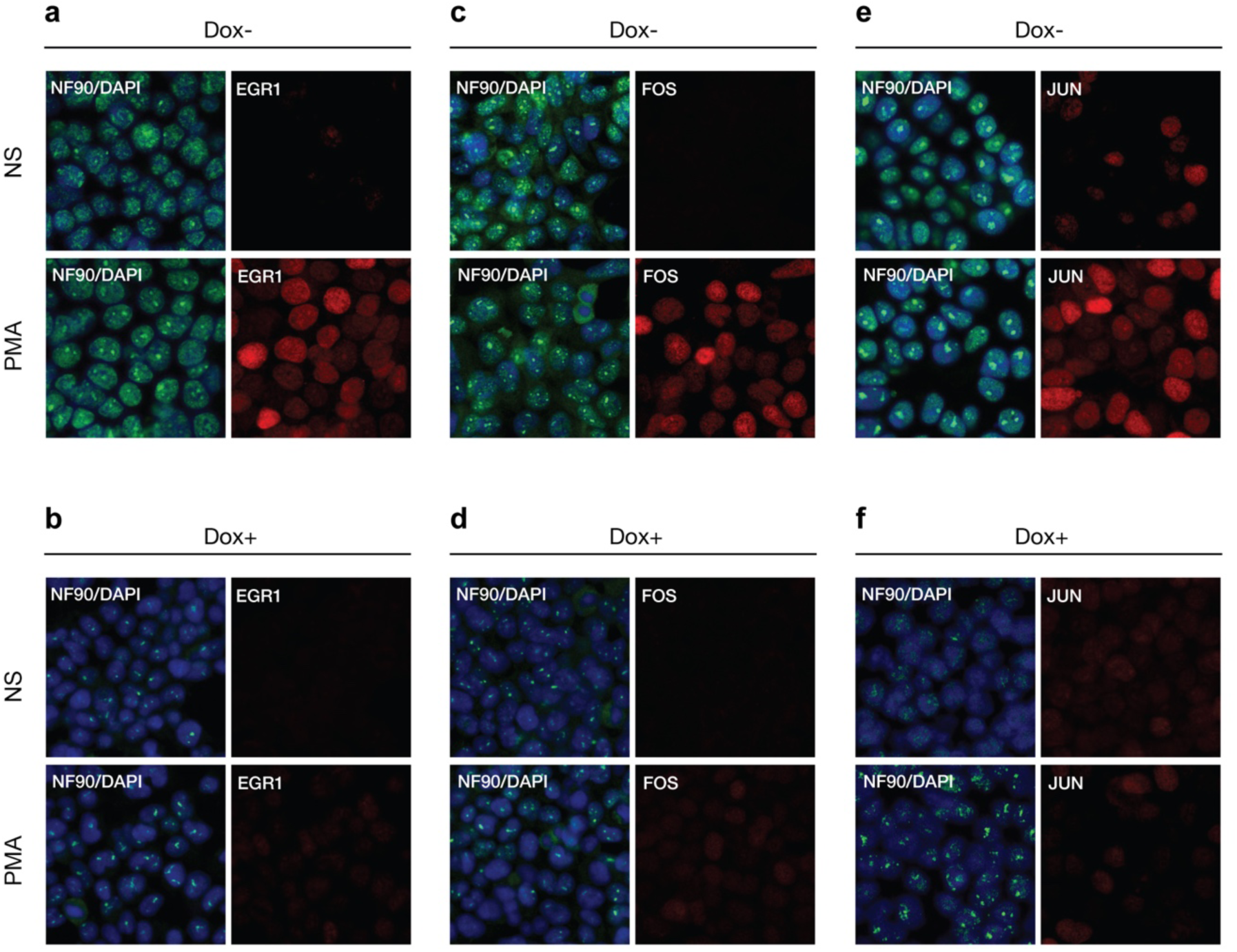
Reduced NF90 expression attenuated inducible expression IEGs by immunofluorescence analysis. HEK293 cells stably transfected with pINDUCER10-shNF90 (D2) or pINDUCER10-shNF45 (D5) were treated without or with doxycycline for 96 h, then seeded on chamber slides, and serum starved for 24 h and stimulated by PMA (20 ng/ml) for 2 h for maximal protein expression of IEG. Slides were incubated with mouse monoclonal antibodies against NF90, and rabbit monoclonal antibodies against *EGR1* (**a-b**), *FOS* (**c-d**), or *JUN* (**e-f**), then incubated with fluorescent conjugated secondary antibodies anti-mouse Alexa Fluor 488 and anti-rabbit Alexa Fluor 594. Slides were counterstained with DAPI, then visualized by confocal fluorescent microscopy. Panels **b,d,f** show doxycycline-mediated shRNA knockdown compared to **a,c,e**, respectively.

**Figure 5.**
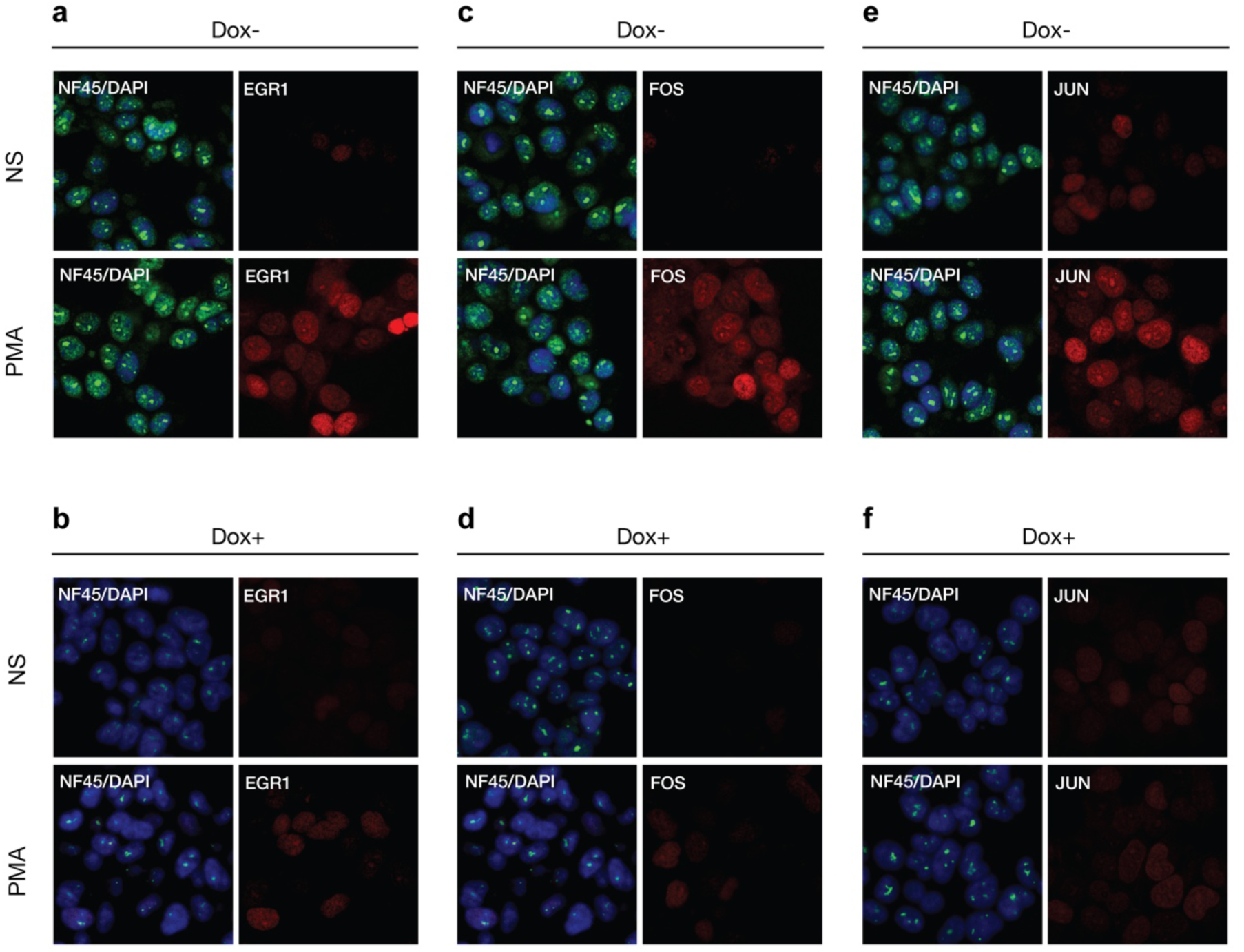
Reduced NF45 expression attenuated inducible expression IEGs by immunofluorescence analysis. HEK293 cells stably transfected with pINDUCER10-shNF90 (D2) or pINDUCER10-shNF45 (D5) were treated without or with doxycycline for 96 h, then seeded on chamber slides, and serum starved for 24 h and stimulated by PMA (20 ng/ml) for 2 h for maximal protein expression of IEG. Slides were incubated with mouse monoclonal antibodies against NF45, and rabbit monoclonal antibodies against *EGR1* (**a-b**), *FOS* (**c-d**), or *JUN* (**e-f**), then incubated with fluorescent conjugated secondary antibodies anti-mouse Alexa Fluor 488 and anti-rabbit Alexa Fluor 594. Slides were counterstained with DAPI, then visualized by confocal fluorescent microscopy. Panels **b,d,f** show doxycycline-mediated shRNA knockdown compared to **a,c,e**, respectively.

HEK293 cells stably expressing doxycycline-regulated shN90 (D2) were cultured on glass chamber slides without or with doxycycline for 96 h before serum starvation overnight followed by stimulation with PMA for 2 h. Cells were fixed with methanol, membranes were permeabilized with Triton X-100 for incubation with primary mouse monoclonal antibody to NF90 together with rabbit monoclonal antibodies to *EGR1*, *FOS*, or *JUN*. All microscope exposure times for a given detection channel wavelength were identical across cell stimulation conditions (**Figure 4**). The NF90 immunoreactivity overlapped with DAPI staining, consistent with nuclear localization of NF90; there was punctate staining within the nuclei consistent with prior reports of NF90 interactions with nucleoli (42). In D2 cells treated with doxycycline for 96 h (Dox+), the immunofluorescence intensity of NF90 was substantially reduced compared to cells not exposed to doxycycline (Dox-) (**Figure 4b,d,f** vs. **a,c,e**). Compared to Dox– nonstimulated (NS) D2 cells, Dox– D2 cells stimulated with PMA (20ng/ml for 2 h), demonstrated clear induction of *EGR1*, *FOS*, or *JUN* proteins within nuclei (**Figure 4a,c,e**; compare PMA vs. NS). In contrast to Dox– D2 cells (**Figure 4a,c,e**), Dox+ D2 cells exhibited marked attenuation of PMA-induced expression of *EGR1*, *FOS*, or *JUN* proteins (**Figure 4b,d,f**; compare PMA vs. NS).

Similarly, HEK293 cells stably expressing doxycycline-regulated shN45 (D5) were visualized using immunofluorescence microscopy (**Figure 5**). The NF45 immunoreactivity overlapped with DAPI staining, consistent with nuclear localization of NF45; there was punctate staining within the nuclei suggestive of NF45 interactions with nucleoli. Compared to Dox– D5 cells (**Figure 5a,c,e**), cells treated with doxycycline for 96 h (Dox+) demonstrated substantial attenuation of NF45 expression (**Figure 5b,d,f**). Stimulation of serum-starved Dox– D5 cells with PMA for 2 h induced substantial expression of *EGR1*, *FOS*, or *JUN* (**Figure 5a,c,e**; compare PMA vs. NS) proteins in nuclei. In contrast, stimulation of Dox+ D5 cells with PMA for 2 h showed marked attenuation of PMA-induced expression of *EGR1*, *FOS*, or *JUN* proteins (**Figure 5b,d,f**; compare PMA vs. NS).

These results from immunofluorescence microscopy demonstrate in single cells that knockdown of NF90 or NF45 proteins are associated with attenuation of PMA-induction of immediate early transcription factors, *EGR1*, *FOS*, and *JUN*.

Taken together, we demonstrate dynamic chromatin occupancy by NF90 and NF45 at the proximal promoters of *EGR1*, *FOS*, and *JUN*. RNAi mediated knockdown of NF90 or NF45 specifically attenuates PMA-inducible RNA and protein expression of immediate early transcription factors *EGR1*, *FOS*, or *JUN*. We propose that NF90 and NF45 are chromatin regulators of IEG transcription.

## DISCUSSION

In this study, we present the first experimental evidence in HEK293 cells that NF90 and NF45 exhibit dynamic chromatin occupancy at the proximal promoters of the IEG transcription factors, *EGR1, FOS* and *JUN*. We also demonstrate their contribution to the inducible gene expression of the immediate early response to cellular stimulation.

In serum-starved, nonstimulated HEK293 cells, NF45 chromatin occupancy was prominent at the proximal promoters of immediate early transcription factors *EGR1*, *FOS* and *JUN*. Upon PMA stimulation, NF45 chromatin occupancy decreased rapidly at 30 minutes, and was further decreased at 60 minutes. Reciprocally, in non-stimulated cells HEK 293 cells, NF90 exhibited modest chromatin occupancy at the proximal promoters of *EGR1*, *FOS* and *JUN*. Upon PMA stimulation, NF90 chromatin occupancy increased rapidly at 30 minutes, and continued to increase at 60 minutes. The kinetics of NF90 and NF45 chromatin association at these proximal promoters is consistent with NF90 and NF45 serving essential roles in regulating the rapid induction of IEGs that occurs on the timescale of 30 minutes to 2 hours.

Both NF45 and NF90 contribute positively to expression of IEG transcription factors *EGR1*, *FOS* and *JUN*. Doxycycline-regulated RNAi knockdown of NF45 or NF90 significantly attenuated PMA-inducible expression of *EGR1*, *FOS* and *JUN* at the levels of mRNA and protein. These results are consistent with literature describing a positive correlation of NF45 and NF90 expression levels with cell growth and proliferation in ESCs (26,27) and diverse cancers (28-33).

The presence of NF45 pre-existing at the proximal promoters of IEGs identify it as a potential pioneer factor capable of interacting with nucleosomal DNA (**Figure 6**, Nonstimulated). The unexpected decrease in NF45 occupancy at proximal promoters of *EGR1*, *FOS* and *JUN* following PMA stimulation might suggest that NF45 operates as a transcriptional repressor. Instead, we propose that NF45 operates as a chromatin remodeler through enzymatic activity intrinsic to NF45 or through recruitment of chromatin remodeling factors. NF45 creates access to promoter DNA sequences for transcriptional activators such as SRF, NF-κB, CREB and NF90. By 30 minutes of PMA stimulation, NF90 is recruited to the proximal promoters of *EGR1*, *FOS* and *JUN* (**Figure 6**, Stimulated, 30 min). In this model, NF45 is positioned as a factor pre-existing at the proximal promoters of rapidly inducible genes, and operates as the principal locus-targeting subunit of the NF90-NF45 gene regulator complex.

**Figure 6.**
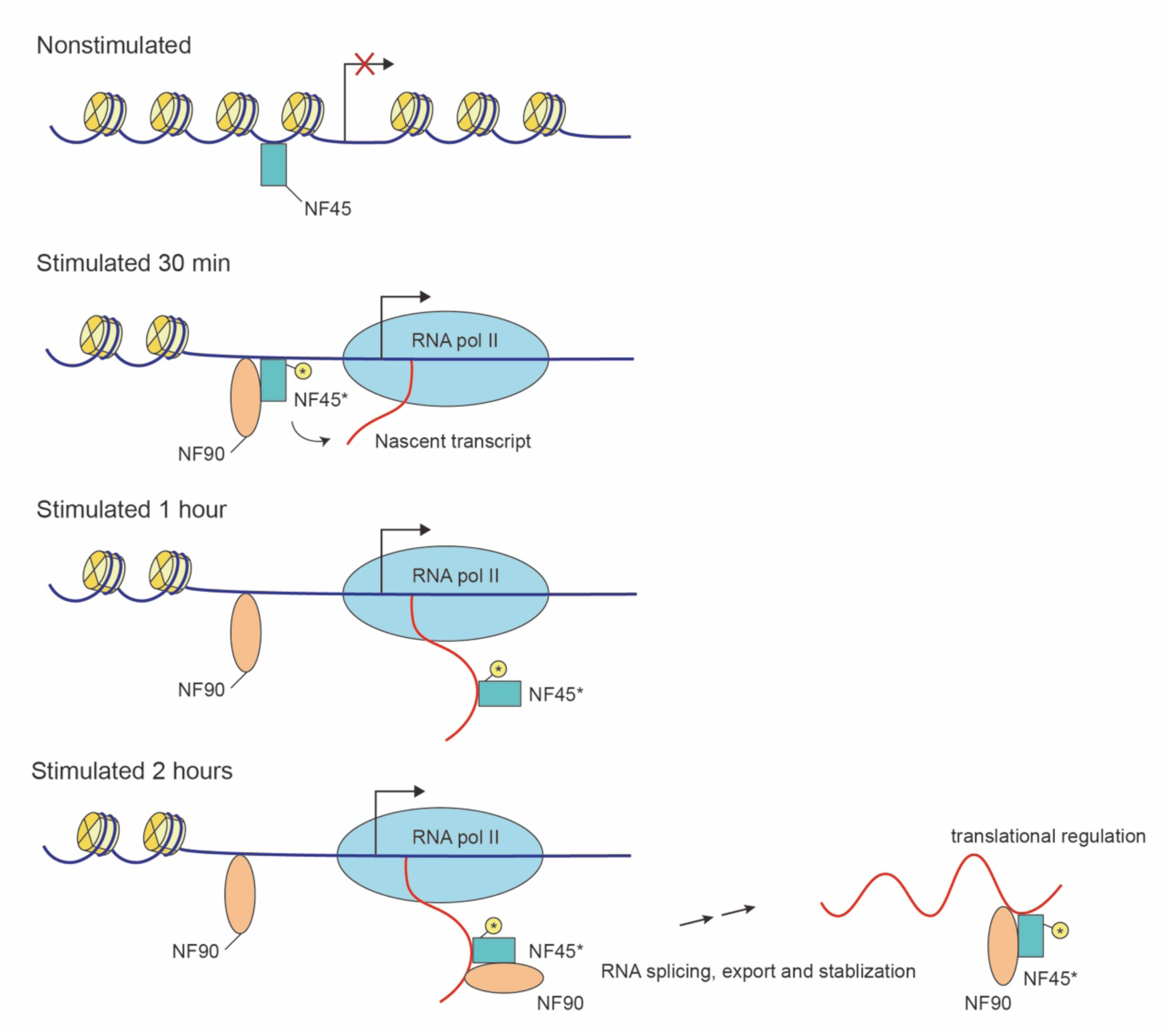
Proposed model for NF90-NF45 regulation of inducible IEG expression upon cellular stimulation. (1) Nonstimulated: Locus is closed. NF45 pre-exists as a nucleosomal DNA-interacting pioneer factor at the proximal promoter of IEGs. (2) Stimulated 30 min: Locus opens. NF45 becomes activated (designated as NF45*, M = methylation) and recruits NF90 to the proximal promoter through shared DZF dimerization domain. RNA polymerase II (RNA Pol II) is recruited and initiates transcription. (3) Stimulated 1 hour: NF45 is released from the proximal promoter and binds to the nascent transcript. (4) Stimulated 2 hours: NF90-NF45 complex binds nascent transcript, and mediates RNA splicing, export, and stabilization, including regulation of translation.

The pre-existing NF45 molecules at the proximal promoters of IEGs are likely targets of signaling initiated at the plasma membrane, such as phorbol ester activation of protein kinase C (PKC) and downstream phosphorylation cascades. Previous proteomics studies have identified NF45 to be diversely modified ^2^, including mono- and di-methylation of the N-terminal arginine/glycine/glycine (RGG) domain, as well as phosphorylation, acetylation, and ubiquitination. The RGG domain has previously been shown to interact with nucleic acids (38). NF110, a splice variant of NF90 that also heterodimerizes with NF45 to form a NF110-NF45 complex, has been shown to be a substrate and regulator of Protein-arginine methyltransferase I (PRMT1) in mammalian cells (39). Methylation of the RGG domain *in vitro* has been shown to inhibit its ability to interact with DNA (38).

The unexpected dynamic and reciprocal chromatin association by NF90 and NF45 is also the first experimental evidence that they are capable of independent interactions at proximal promoters of immediate early transcription factors, in contrast to frequent observations of NF90 and NF45 as a heterodimer. The heterodimerization between NF90 and NF45 is mediated through their shared DZF domains (17), and this interaction may be regulated by cell signaling to rapidly recruit NF90 to the NF45 pre-associated at the proximal promoters of IEGs.

The reduction in NF45 chromatin occupancy at the proximal promoters at 30 and 60 minutes of stimulation is consistent with a model in which NF45 is released from chromatin and becomes available to bind to RNA nascently transcribed from this locus, serving to chaperone RNA through splicing (40,41), nuclear export, stabilization (42,43) and translational regulation (44,45) (49) (**Figure 6**). Nakadai et al. (22) were the first to propose that NF45 and NF90 operate as transcriptional coactivators of *FOS*, and are transferred from the promoter to the RNA Pol II complex during elongation and ultimately bind mRNA to facilitate export into the cytoplasm. In our model, stimulation initiated at the cell membrane transmits signals to NF45 pre-associated at the proximal promoters of *EGR1*, *FOS* and *JUN,* inducing posttranslational and/or conformational changes in NF45 that rapidly recruit NF90 to chromatin and facilitate release of NF45 from chromatin (**Figure 6**, NS, 30, 60 mins).

Targeted disruption of NF90 in mice exhibited perinatal lethality and stunted growth (19). Targeted disruption of NF45 in mice exhibited embryonic lethality at the blastocyst stage^3^. In this study, we demonstrate that RNAi-mediated knockdown of either NF90 or NF45 attenuated inducible expression of immediate early transcription factors, *EGR1*, *FOS* and *JUN*. These data underscore the dual requirement of NF90 and NF45 for IEG expression. NF90 and NF45 regulate gene expression at the levels of chromatin occupancy, transcription, RNA splicing, nuclear export, stabilization and translation.

The authors declare that they have no conflicts of interest with the contents of this article.

## Acknowledgements

We thank Jessica Wu for help with illustrating the proposed model. National Institutes of Health Grant NIAID 1-R01-AI39624 provided support for PNK and LF.

## Footnotes

1 Zhao, G., Wu, T.H., Shi, L., Kao, P.N. manuscript in preparation.

2 https://www.phosphosite.org/proteinAction.action?id=9766&showAllSites=true

3 Zhao, G., Wu, T.H., Shi, L., Kao, P.N. manuscript in preparation.

